# The glutamine-aspartate metabolic tradeoff between fast growth and metabolic flexibility

**DOI:** 10.1101/2025.05.12.652188

**Authors:** Karin Ortmayr, Meret Ringwald, Laurentz Schuhknecht, Sébastien Dubuis, Eleni Paunossis, Martina Blaesi, Charly Jehanno, Sarah D. Grieder, Mohamed Bentires-Alj, Moritz Mall, Mattia Zampieri

**Affiliations:** Department of Biomedicine, University of Basel, Basel, Switzerland; Institute of Molecular Systems Biology, ETH Zürich, Switzerland; Department of Surgery, University Hospital Basel, Basel, Switzerland; Department of Pharmaceutical Sciences, Faculty of Life Sciences, University of Vienna, Austria; Cell Fate Engineering and Disease Modeling Group, German Cancer Research Center (DKFZ) and DKFZ-ZMBH Alliance, 69120 Heidelberg, Germany; HITBR Hector Institute for Translational Brain Research gGmbH, 69120 Heidelberg, Germany; Central Institute of Mental Health, Medical Faculty Mannheim, Heidelberg University, 68159 Mannheim, Germany

## Abstract

Cancer cells develop unique addictions to nutrients, offering attractive opportunities for clinical intervention. However, how cancer cells orchestrate and benefit from metabolic rewiring remain fundamental open questions. Here, we developed a systematic framework to investigate nutrient dependencies and applied it to study glutamine addiction. By measuring metabolite exchange rates across 54 cancer cell lines with different levels of dependency on extracellular glutamine and in human stem cells, we discovered that a dualistic regulation of glutamate biosynthesis from glutamine or aspartate, driven by the tissue of origin, generates a tradeoff between fast growth and metabolic flexibility. Monitoring dynamic metabolic adaptation to glutamine deprivation showed that the role of glutamine as limiting substrate in purine biosynthesis is not responsible for glutamine dependency. Rather, growth arrest is mediated by glutamine as a signaling molecule. Our approach opens new opportunities to map, mechanistically investigate and pharmacologically interfere with metabolic dependencies, beyond glutamine addiction.

## Introduction

Reprogramming of metabolism^1,2^ allows cancer cells to sustain rapid growth, maintain redox homeostasis, avoid immune response and adapt to new environmental niches^3–6^. A unique characteristic of the widespread metabolic reprogramming in cancer cells is the addiction to nutrients that are not only consumed at a much higher rate than in normal healthy cells, but also become essential for cancer cell survival, proliferation and metastatic potential^7 8^. Metabolic specialization in cancer creates unique opportunities in diagnosis and for selective cancer therapies^9–11^. However, despite an ever-increasing discovery of cancer nutrient addictions^12,13^ and their potential clinical impact^14^, a systematic understanding of how cancer cells regulate and coordinate nutrient uptake and metabolism to sustain cell proliferation and survival remains challenging.

One of the most studied nutrient dependency in cancer is glucose addiction^15,16^. Not only do cancer cells exhibit high glucose uptake, but unlike healthy cells, cancer cells ferment glucose to lactate even when oxygen is not limiting. A phenomenon known as the Warburg effect^15^. Since the discovery of the Warburg effect, many more nutrient dependencies in cancer cells have been identified ^14,17–21^. Several involve nominally non-essential metabolites that can be synthesized from other nutrients, like amino acids, fatty acids or acetate ^22^. For most, it remains unclear whether the nutrient dependency across different tumor types has similar functional properties^23 24^. The major difficulty in studying these phenomena is the complexity and diversity with which cells can interchangeably utilize different nutrients (i.e., metabolic flexibility) or use the same nutrient to fulfill different purposes – e.g., catabolism, anabolism or signaling (i.e., metabolic plasticity) ^6^.

An emblematic example is glutamine addiction^25^. Glutamine is a highly versatile nutrient that can participate in many essential cellular functions, such as energy formation- as a major TCA anaplerotic substrate, redox homeostasis - as substrate for glutathione, macromolecular synthesis - as a precursor of other amino acids, fatty acids, nucleotides, and signaling - as a key allosteric regulator of mTOR^26^. Moreover, glutamine is one of the most abundant amino acids in plasma. The high demand for glutamine in proliferating cancer cells *in vitro* ^27 28 29,30^ has now also been shown *in vivo*^31 32 33^, where it can play a key role in mediating metastasis^34^. However, the mechanisms by which extracellular glutamine limitation prevents growth of some cancer cells but not others, and whether the same functional role of glutamine drives addiction across different cancer types, remain unclear. Even *in vitro*, where the dependency on extracellular glutamine is aggravated by high cystine concentrations in the media^35 36^, a systematic characterization of essentiality of extracellular glutamine in largely diverse cell types, its underlying origin, and possible bypass mechanisms is still missing. Here, we present a systematic approach to shed mechanistic insights on nutrient dependencies in cancer. We applied it to study extracellular glutamine dependency across a panel of 54 largely diverse cancer cell lines and in human embryonic stem cells (hESC) cells differentiating into neurons. By using multivariate statistical analysis, we combined high-throughput measurements of nutrient uptake and byproduct secretion rates with automated time-lapse microscopy monitoring the phenotypic response of cells to glutamine deprivation (Figure 1A). We show that the universal role of glutamine in limiting growth rate by limiting purine biosynthesis alone cannot explain glutamine addiction. Instead, growth arrest upon extracellular glutamine deprivation is the result of glutamine’s role as a signaling molecule rather than as a limiting metabolic substrate. Notably, we revealed a mechanism largely driven by the tissue of origin by which a dualistic regulation of aspartate and glutamine metabolism enables cells to become independent from extracellular glutamine. Understanding the mechanisms that allow to bypass glutamine dependency can open new translational opportunities in rational design of combination therapies and reveal fundamental principles in adaptation and transformation of cancer cells.

**Figure 1.**
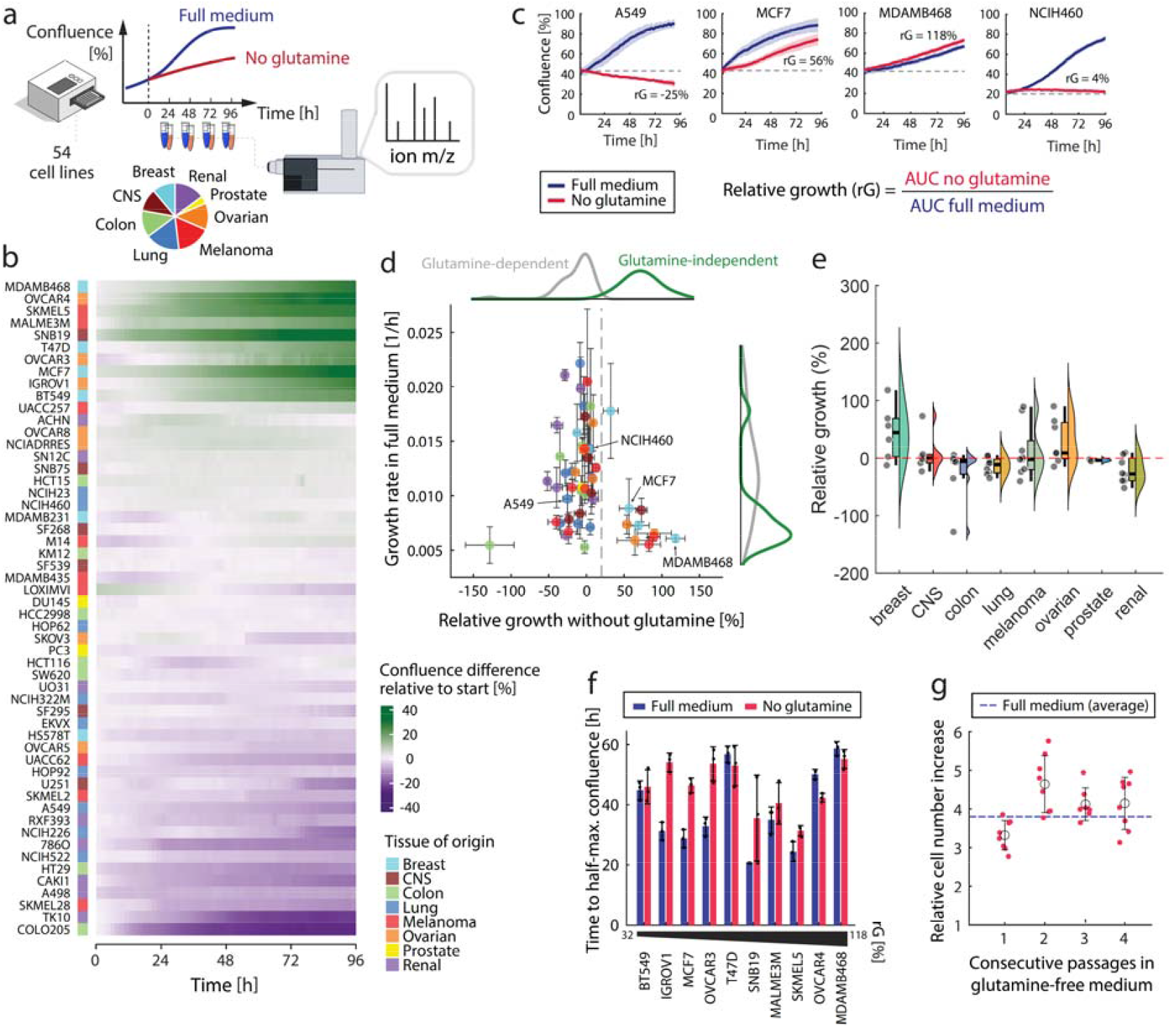
Phenotypic characterization of 54 cell lines from the NCI-60 panel in response to glutamine deprivation. **(a)** Schematic overview of the experimental workflow and cell lineages. Cell confluence was measured using automated brightfield microscopy in 1.5-hours intervals for 96 hours and (exo-) metabolomics samples were collected every 24 hours. **(b)** Cell confluence up to 96 hours after glutamine removal from the medium. For each cell line, the increase in confluence over time with respect to confluence before media change is reported. **(c)** Example growth curves for MDAMB468, NCHI460 and A549 in either RPMI-1640 medium with 2 mM glutamine (blue) or RPMI-1640 medium lacking glutamine (red). The ability to grow without glutamine was estimated from the growth data using the relative growth index (rG). The dashed grey line indicates the starting confluence and reference for calculating the area under each growth curve (AUC). **(d)** Comparison of growth rate in full medium (containing glutamine) versus relative growth in glutamine-free medium. Data are mean ± standard deviation across three replicates. Marginal distributions of growth rate and relative growth in glutamine-free medium are shown for glutamine-dependent cell lines (< 20% relative growth, grey) and glutamine-independent cell lines (≥ 20% relative growth, green). **(e)** Relative growth of cell lines grouped by their tissue of origin **(f)** Estimated growth dynamics at the transition from full- to glutamine-free medium for 10 glutamine-independent cell lines (relative growth ≥ 20%). Data shown is the time until 50% of the maximum confluence was reached in either RPMI-1640 containing 2 mM glutamine (full medium, blue) or glutamine-free medium (red), as mean across three replicates and with error bars indicating the standard deviation. **(g)** Relative increase in cell number over 4 consecutive passages of MDA-MB-468 cells in glutamine-free medium. A defined number of cells was seeded in T25 cell culture flasks and maintained in glutamine-free medium. After 4 days, cells were counted (= 1 passage) using an automated cell counter and re-seeded at the same cell density in glutamine-free medium (= second passage). Data points represent cell counting results from two replicate cultures and 4 technical replicates (cell counting chambers). Black circles and error bars indicate the mean and standard deviation, respectively.

## Results

### Inherent variability in glutamine dependency across cancer cell lines

To better understand glutamine addiction in cancer cells, we selected 54 adherent cancer cell lines from 8 different tissues (NCI60 cancer cell line panel)^37^ and used time-lapse microscopy to systematically investigate their inherent ability to grow upon acute deprivation of extracellular glutamine. Specifically, cells were seeded in 96-well plates in RPMI-1640 medium (i.e. supplemented with dialyzed fetal bovine serum, 2 g/L glucose and 2 mM glutamine) and grown for 48 hours before the medium was exchanged to RPMI-1640 medium with or without glutamine (Figure 1a and Supplementary Figure 1). We subsequently monitored cell confluence every 1.5 hours for 96 hours (Figure 1b) to estimate the ability of cells to grow without glutamine using the relative-growth index, defined as the ratio of the area under the growth curve of cells growing without and with glutamine (Figure 1c, Supplementary Information 1 and Supplementary Figure 2). While 0 or negative relative-growth values indicate growth arrest or cell detachment, cell lines with a relative growth close to 1 exhibit comparable growth with and without glutamine (Figure 1b-c). We observed a separation of the cell lines into two groups with significantly different growth dynamics upon glutamine limitation (Figure 1d). 10 cell lines were able to continue growing in medium lacking glutamine, while of the remaining 44 cell lines 31 immediately halted growth and 13 even exhibited a progressive reduction in confluence (Figure 1b, relative growth < 0), indicative of cell detachment and/or death (Supplementary Information 1). While cell lines of the same tissue type can exhibit largely different levels of glutamine dependency (Figure 1d), ovarian and breast cancer cell lines exhibited the highest fraction of glutamine-independent cell lines (Figure 1e). Overall, we found that cells able to grow without extracellular glutamine supplementation showed, on average, lower growth rates already in full medium than glutamine-dependent cell lines (Figure 1d). Strikingly, all 10 cell lines considered as glutamine-independent according to the relative growth index (Figure 1e, p-value<0.01) grew uninterruptedly upon glutamine deprivation (Figure 1f-g). This ability to promptly grow after medium exchange suggested that the metabolic state before glutamine deprivation determines whether cells can sustain growth without glutamine.

### Aspartate uptake is increased in glutamine-independent cell lines

Because the 10 glutamine-independent cell lines resumed growth with almost no delay upon extracellular glutamine deprivation, we hypothesized that a different metabolism and usage of available nutrients even before glutamine deprivation is what distinguishes glutamine-dependent from -independent cell lines (Figure 1). To test our hypothesis, we measured relative differences in nutrient uptake and secretion rates of metabolic byproducts in all 54 cell lines before glutamine deprivation (i.e. in full medium containing glutamine). To that end, we used high-throughput non-targeted mass spectrometry (FIA-TOFMS^38^) to measure metabolite levels in cell culture supernatants at several time-points during growth (Figure 1a). Based on the exometabolome measurements, we calculated exchange rates (ion intensity per cell volume and time, a.u./V/h) for 2‘443 putatively annotated metabolites and combined these new measurements with previously determined estimates of glucose uptake and lactate secretion rates^39^ (Supplementary Information 2). It is worth noting that, unlike previous absolute estimates of metabolite exchange rates that measured only end-point concentrations at approximately 80% confluence^40^, our relative estimates are based on multiple measurements every 24 hours over four days of growth (Figure 1a and 2a) and account for differences in cell volumes between cell lines^39 41 42 43^.

We detected 37 of the 42 components in the chemically defined basal RPMI-1640 medium (i.e. amino acids, vitamins, co-factors, Supplementary Information 2). Differences in the exchange rates of the media components across the 54 cell lines followed two main trends that were largely tissue-independent (Figure 2b). Several amino acids including glutamine, cystine and essential amino acids^44^ histidine, (iso)leucine, lysine, methionine, phenylalanine, threonine, tryptophan and valine exhibited uptake rates correlating with glucose consumption. The exchange rates of mostly non-essential amino acids, like glutamate, proline and arginine (synthesized from α-ketoglutarate) or aspartate and asparagine (both derived from oxaloacetate), together with nucleotide intermediates, such as thymidine, hypoxanthine, and cofactors, like folate, biotin and GSH followed a different trend which largely anticorrelated with glucose – i.e. the larger the uptake of glucose and essential amino acids, the lower the uptake of non-essential amino acids and nucleotide intermediates. Such a global correlative trend suggests highly coordinated strategies in the uptake and utilization of essential and non-essential nutrients. Besides components of the chemically defined basal medium, we detected other putative extracellular metabolites, likely originating from the dialyzed fetal bovine serum in the complete growth medium, or the secretion of diverse metabolic products (e.g. N2-Formyl-N1-(5-phospho-D-ribosyl)glycinamide, m/z 314.051). The exchange rates followed the two major trends observed for the main medium components (Figure 2c) and reproduced well-known relationships between nutrient uptake rates and secretion of metabolic byproducts, such as the correlation between glutamine uptake and glutamate secretion^40^ (Spearman R -0.54, Figure 2d), or the anti-correlation between glutamate secretion and cystine uptake (Spearman R -0.61, Figure 2e), reflecting the cystine antiport mechanism^45^.

**Figure 2.**
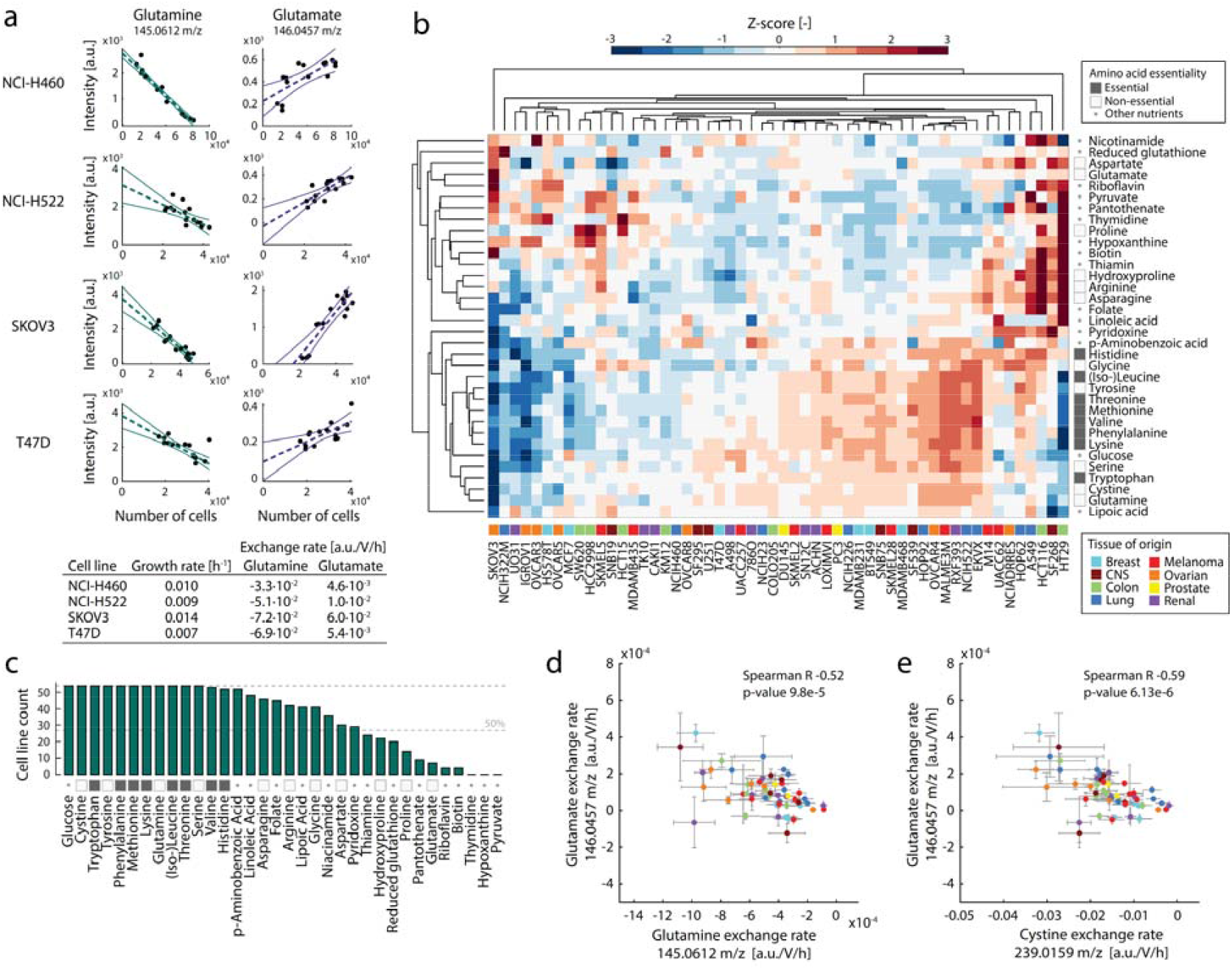
Exometabolome profiling of 54 adherent cancer cell lines in RPMI-1640 medium with 2 mM glutamine. **(a)** Estimation of exchange rates (a.u./V/h) from measured relative ion abundances using least-squares fitting analysis of ion intensity as a function of cell numbers. Solid lines represent the 95% confidence interval. The optimal fitted slope is then corrected for characteristic cell volume and growth rate previously estimated for each cell line^46^. Dashed lines represent the optimal fit based on a linear regression scheme and solid lines delimit the zone of 95% confidence. **(b)** Heatmap of z-scored normalized uptake/secretion rates for the RPMI-1640 medium components annotated in our metabolome dataset. Colored boxes indicate the tissue of origin for each cell line. For amino acids, essentiality is indicated in gray or white boxes. **(c)** Consumption of major nutrients by 54 different cell lines. For each RPMI medium component, we report the number of cells with a net consumption. For amino acids, essentiality is indicated in gray or white boxes (legend see panel b). **(d-e)** Correlation between exchange rates of glutamine and glutamate (panel d) and glutamate and cystine (panel e) across 54 adherent cancer cell lines grown in RPMI-1640 medium with 2 mM glutamine.

Glutamine-independent cell lines exhibited on average a slower growth rate in glutamine-containing medium (Figure 1d) and showed only minor lag times or growth reduction upon extracellular glutamine deprivation (Figure 1e). Therefore, we hypothesized that glutamine-independent cell lines may inherently have lower consumption rates of glutamine with respect to glutamine-dependent cell lines. However, we found that glutamine uptake rates were not significantly different between the two groups (q-value = 0.8, Figure 3a-b). Instead, out of the 2‘443 measured putatively annotated metabolites, aspartate exhibited a unique and most significant difference (q-value = 0.019, Storey correction) (Figure 3a-b). While all glutamine-independent cell lines featured a net uptake of aspartate, glutamine-dependent cell lines exhibited lower uptake or a net secretion of aspartate (Figure 3b). Notably, the expression level of the aspartate transporters EAAT1 and EAAT2 do not correlate with aspartate exchange rates (i.e. Spearman correlation of 0.08 and 0.3, respectively).

**Figure 3.**
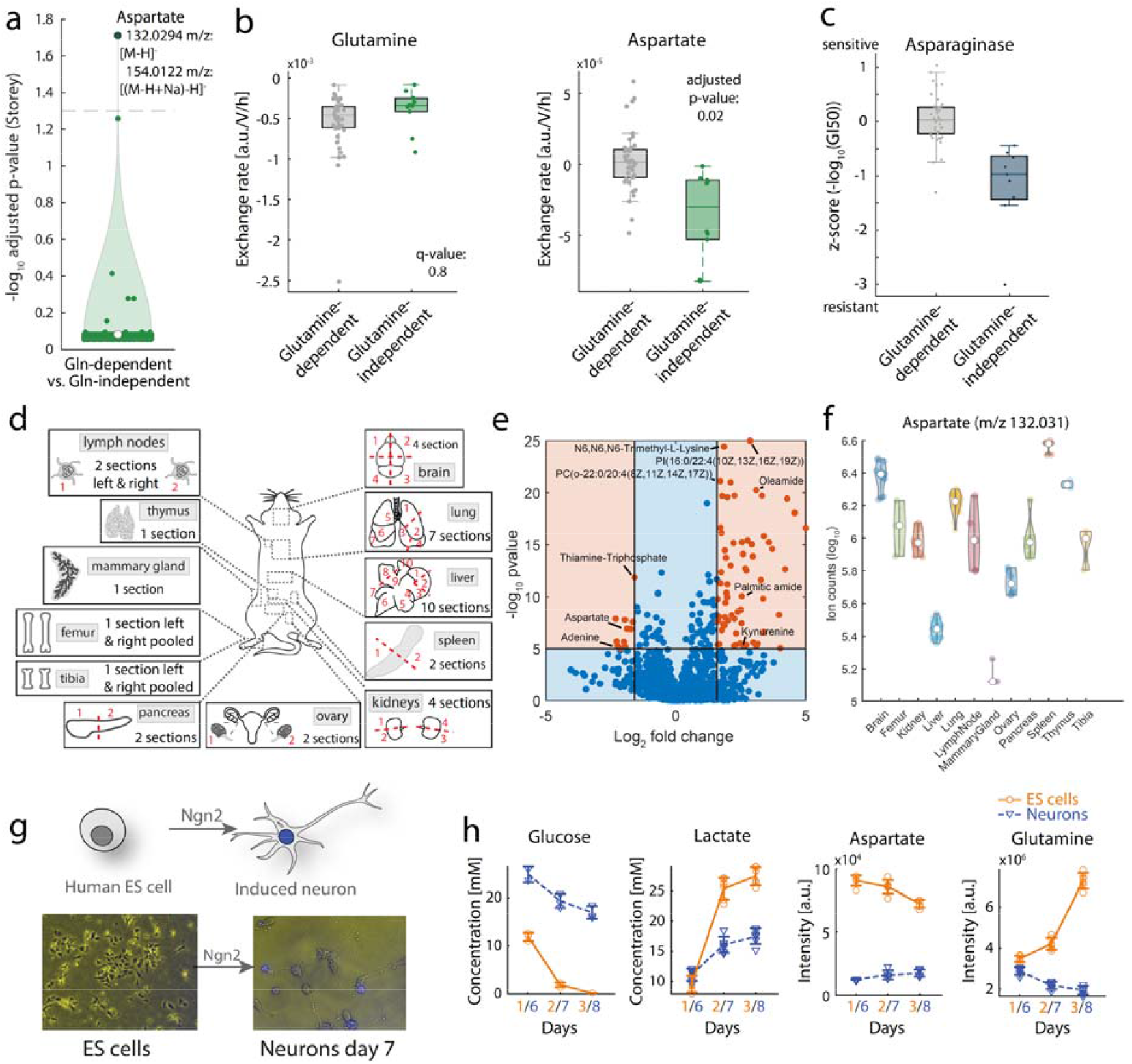
Aspartate and glutamine metabolism in glutamine in/dependent cell lines and during hESC differentiation. **(a)** Comparison of uptake/secretion rates between glutamine independent and dependent cell lines (Student’s t-test, Bonferroni correction for multiple hypothesis testing). **(b)** Comparison of glutamine and aspartate exchange rates in glutamine-independent (grey) and dependent cell lines (green). **(c)** Comparison of sensitivity (IC50-value) to asparaginase treatment in glutamine independent (grey) and dependent cell lines (green). **(d)** Schematics of organ sites of FVB mice sampled for metabolome profiling. **(e)** Volcano plot showing differential abundance analysis of metabolites from FIA-TOF analysis of average metabolite levels from all tissue sections in brain, lungs, kidneys *vs* average levels in mammary glands and ovaries. P-values were calculated by a t-test analysis. The horizontal line indicates a p-value of 1e-5. Orange dots represent metabolites with significant differences between the two groups of organ tissues. **(f)** Distribution of aspartate levels across different mouse tissues. **(g)** Schematic and microscopic images of the experimental setup for profiling metabolic changes during differentiation from embryonic stem cells (ES) to neurons. **(h)** Glucose, lactate, aspartate and glutamine concentrations in the media supernatant of pluripotent stem cells (day 1-2-3) and neurons (days 6-7-8 after transcription factor-mediated differentiation).

Moreover, by comparing the sensitivity to 21’738 drugs across the 54 cell lines^47^, we found that asparaginase showed the most significant differential sensitivity between glutamine-independent and - dependent cell lines (q-value = 2.17e-04), in that glutamine-independent cell lines were on average less sensitive to asparaginase treatment (i.e. higher GI_50_) (Figure 3c) (Supplementary Information 5). Asparaginase is a therapeutic enzyme that converts asparagine to aspartate in the extracellular space, thereby depleting asparagine and forcing cells to synthesize asparagine from aspartate. The decreased sensitivity of glutamine-independent cell lines to asparaginase activity is consistent with an increased aspartate uptake and therefore substrate availability for asparagine biosynthesis. Moreover, the differential sensitivity of glutamine-dependent *vs* independent cell lines to asparaginase strengthen the role of aspartate in bypassing glutamine addiction. The observed increased aspartate uptake in glutamine-independent cell lines agrees with earlier reports, which showed that higher availability of cytosolic aspartate can improve cell survival under glutamine deprivation^48 49^. We therefore hypothesized that increasing aspartate uptake could be part of a common strategy to adapt to glutamine limitation that is shared across widely different cell types beyond cancer cells.

### Tissue of origin and differentiation dictates aspartate metabolism and glutamine dependency

Ovarian and breast cancer cell lines exhibited a higher fraction of glutamine-independent cell lines (Figure 1e) compared to other tissue-derived cancer cell lines (i.e. 54% *vs* 7%), suggesting that glutamine dependency and preference for consumption over secretion of aspartate may be driven by the tissue of origin. To address this question in an *in vivo* context, we measured levels of 1‘257 putatively annotated metabolites in 12 different mouse organs of FVB mice (Figure 3d) (Supplementary Information 4 and supplementary figure 3). We hypothesized that, because metabolome composition of different organs in mice can reveal organ-specific metabolic patterns that are also relevant to human physiology^50–53^, organs in which cells mainly secrete aspartate would exhibit higher levels of aspartate than tissues in which aspartate is taken up and metabolized, and hence that tissue aspartate levels could report on glutamine dependency in human cancer cell lines. To test our hypothesis, we compared average metabolite levels in organs from which tested NCI60 human cancer cell lines mainly exhibited glutamine dependency (e.g. brain, lung and kidneys), and organs from which human cancer cell lines exhibited glutamine independency (e.g. mammary gland and ovary) (Figure 3d). Remarkably, aspartate was among the metabolites with the most significant (p-value<1e-5) difference between the 2 tissue groups, with almost 4-fold lower levels in mammary glands and ovaries (Figure 3d). To test whether mouse tissue levels of aspartate are predictive of glutamine independence in human cancer cell lines, we investigated if cancer cell lines different from those tested here (i.e. NCI60 cell lines) and originating from tissues with low or high aspartate levels in mouse would exhibit glutamine in-/dependency, respectively. Among the profiled organ tissues, only in the liver aspartate levels were lower than in mammary glands or ovaries (Figure 3f). Notably, consistent with our hypothesis, 9 (SK-Hep1, PLC/PF5, HepG2, HUH6, Hep3B, SNU387, JHH1, HUH7) out of 12 (SNU449, SNU398, MHCC97H) (75%) liver cell lines previously tested for glutamine dependency^54^ were able to grow without supplementing glutamine to the medium (i.e. glutamine independent). Conversely, the pancreas features relatively high levels of aspartate in mice (Figure 3f). Out of 8 pancreatic cell lines^55^ (8988T, Tu8902, Panc1, Miapaca2, PL45, MPanc96, Primary PDAC 1 and 2) tested for glutamine dependency^55^, none were shown to be able to survive without extracellular glutamine (i.e. glutamine dependent). Similarly, high aspartate levels were detected in mouse organs like spleen and the thymus, lymphoid organs that contain various cell lines crucial for immune functions. Consistent with high tissue levels of aspartate, several are the studies showing that immune cells activation requires extracellular glutamine ^56–59^(C57BL/6J splenocytes). Altogether, our results are consistent with previous findings^60^ and indicate that the tissue context can largely define cancer regulation of specific metabolic pathways and nutrient dependencies.

However, whether and how such dependencies emerge during cellular differentiation to a mature specialized tissue type remains an open question. Because independence from exogenous glutamine was reported to occur in pluripotent stem cells^61^, we next tested whether differential uptake of aspartate could also be observed during stem cell differentiation (Figure 3g). We monitored extracellular (Figure 3h) and intracellular (Supplementary Figure 3) metabolic changes in human embryonic stem cells (hESCs) during transcription factor-mediated differentiation into excitatory induced neurons. Consistent with the above findings in tumor cells, we found significant progressive metabolic rewiring in aspartate metabolism (see Supplementary Figure 3 and Supplementary Information 4). Specifically, we observed that while hESC cells exhibit a net consumption of aspartate and secretion of glutamine (Figure 3h), induced neurons exhibited an opposite behavior with a net glutamine consumption and secretion of aspartate (Supplementary Information 4). Hence, our results suggest that differential uptake of aspartate is a fundamental mechanism harnessed also by pluripotent stem cells to proliferate without the need for extracellular glutamine^61^.

### Glutamine-independent cell lines use aspartate to synthesize glutamate

While increased aspartate uptake was a common feature among glutamine-independent cell lines, we observed that 20 of the glutamine-dependent cell lines also showed net uptake of aspartate. This observation could indicate that high aspartate intake is necessary but not sufficient to provide independence from extracellular glutamine. To directly test whether higher intracellular availability of aspartate can be sufficient to bypass glutamine dependency, we sought to increase the cytosolic aspartate levels in glutamine-dependent cell lines using aspartate dimethyl ester to facilitate aspartate diffusion across the cell membrane. We selected four cell lines that differ in aspartate uptake and relative growth upon glutamine deprivation, i.e. the glutamine-independent cell line MDA-MB-468 (net uptake of aspartate), and the three glutamine-dependent cell lines A549 (net secretion of aspartate), SNB75 (with an aspartate exchange rate of 0), and UO31 (net uptake of aspartate). All cell lines were cultivated in full medium with up to 6 mM aspartate dimethyl ester. However, despite a significant (p-value < 0.05) increase in the levels of intracellular aspartate already at 1 mM (Supplementary Figure 4), supplementing dimethyl-aspartate in glutamine-free medium was not enough to resolve glutamine dependency (Figure 4a), suggesting that additional limitations can hinder growth under glutamine limitation. Hence, we investigated the differences in intracellular metabolism between cells that are independent and dependent on extracellular glutamine.

**Figure 4.**
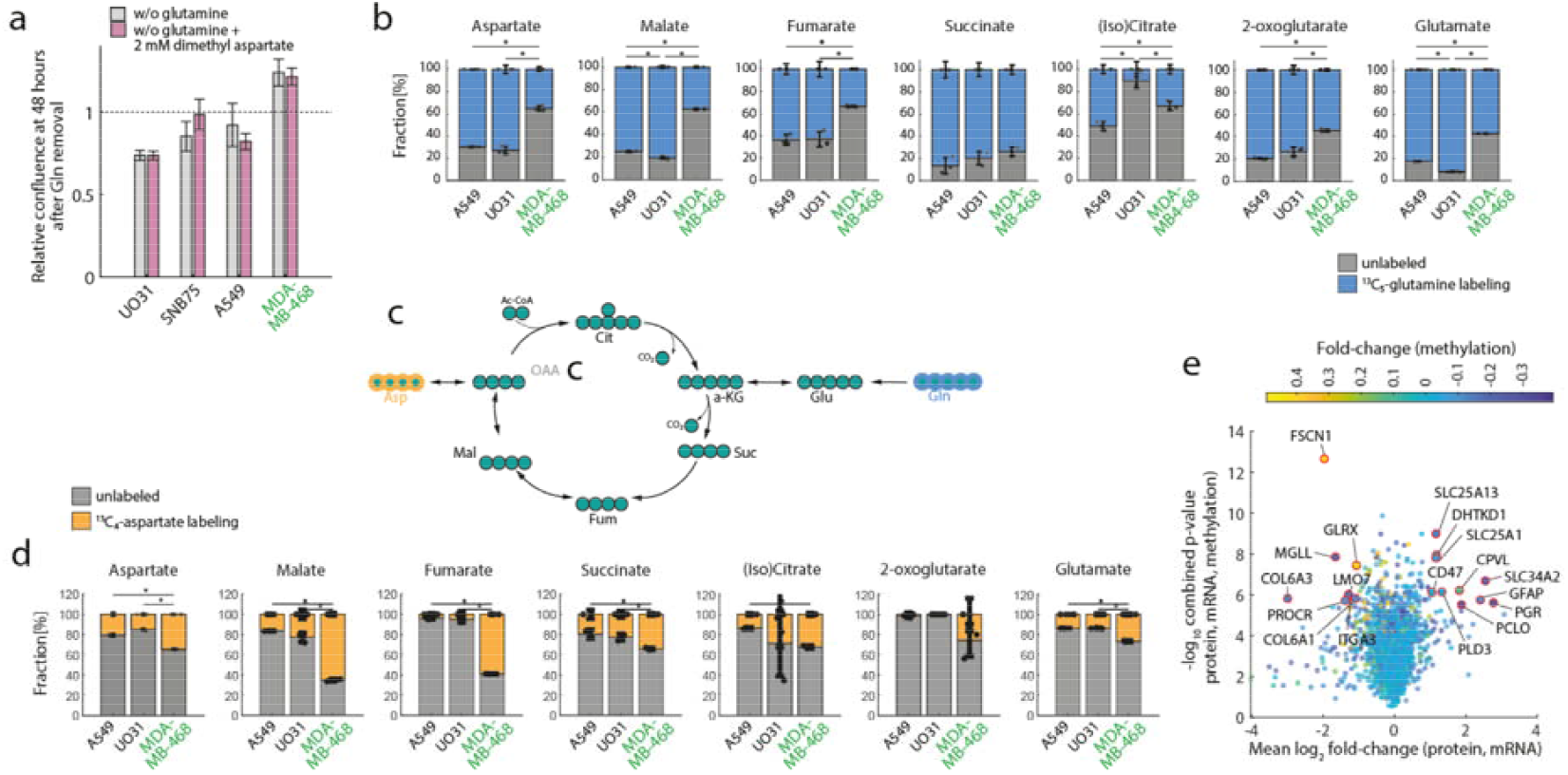
Aspartate and glutamine metabolism in glutamine in/dependent cell lines. **(a)** Three glutamine-dependent (A549, UO31 and SNB75) and one glutamine-independent cell line (MDA-MB-468, in green) were grown in RPMI-1640 medium lacking glutamine (grey) and supplemented with 2 mM aspartate dimethyl ester (red). The confluence 48 hours after glutamine deprivation is divided by the confluence before glutamine deprivation and reported as a ratio (i.e., relative confluence). **(b)** Percent labeling from ^13^C_5_-glutamine (% of total pool) in intracellular aspartate and glutamate in A549, UO31 and MDA-MB-468 grown in RPMI-1640 medium with 2 mM ^13^C_5_-glutamine for 24 h. All isotopologues are lumped together (blue). Data represent the mean ± SEM of 3 biological replicates. **(c)** Schematic of TCA cycle and how aspartate and glutamine respectively can be used as anaplerotic substrates. **(d)** Percent labeling from ^13^C_4_-aspartate (% of total pool) in intracellular aspartate and glutamate in A549, UO31 and MDA-MB-468 grown in RPMI-1640 medium (2 mM glutamine, 150 µM aspartate) supplemented with 150 µM ^13^C_4_-aspartate for 24 h. All isotopologues are lumped together (orange). Data represent the mean ± SEM of 3 biological replicates. **(e)** Average mRNA^62^ and protein_63_ fold changes vs the respective geometric mean of the t-test pvalues calculated for 3039 genes between glutamine independent vs dependent cells. Color code is proportional to fold change in gene methylation ^62^ between glutamine independent vs dependent cells.

To analyze differences in aspartate and glutamine metabolism, we employed isotopic tracing using cell culture media with either ^13^C_5_ -labeled glutamine or with ^13^C_4_-labeled aspartate (50% of the total pool). We selected three cell lines that differ in aspartate uptake and relative growth upon glutamine deprivation, i.e., A549 (glutamine-dependent, net aspartate secretion), MDA-MB-468 (glutamine-independent, net aspartate uptake), and UO31 (glutamine-dependent, net aspartate uptake). Using mass spectrometry, we traced the fates of ^13^C_5_ -glutamine and ^13^C_4_ -aspartate, i.e. the incorporation of the isotopic label in other intracellular metabolites (Figure 4b-d, Supplementary Information 3). Consistent with the differences in aspartate uptake (Figure 3b), the glutamine-independent cell line MDA-MB-468 derived most (>65%) of the aspartate from the medium, while in the two glutamine-dependent cell lines UO31 and A549, aspartate mainly (>70%) originated from the supplemented glutamine (Figure 4b). Similarly, we found that the contribution of glutamine-derived carbon (Figure 4b) into key downstream metabolic products like glutamate, aspartate or fumarate, was significantly (p-value<0.01) lower in the glutamine-independent cell line MDA-MB-468 than in the two glutamine-dependent cell lines (Figure 4b). Notably, analysis of the labeling patterns from ^13^C_4_ -labeled aspartate showed that aspartate contributes to the synthesis of glutamate in glutamine-independent MDA-MB-468 cells (Figure 4d), suggesting that aspartate may alleviate the need for extracellular glutamine. In addition, we found that glutaminase (GLS, converting glutamine into glutamate) expression^62^ in full medium is on average lower in glutamine-independent cell lines, while glutamine synthetase expression (GLUL, converting glutamate into glutamine) is higher (Supplementary Figure 5, Supplementary Information 5). This observation further emphasizes the differential basal regulation of glutamate biosynthesis from glutamine and its role in mediating (in)dependency from extracellular glutamine.

To identify potential regulatory events driving differential metabolism of aspartate and glutamine, we sought differentially regulated genes in glutamine-dependent versus -independent cell lines by using previously published DNA methylome, transcriptome and proteome data for the NCI-60 cancer cell line panel ^62,63^. Among the 249 genes with a significant (p-value<1e-5) and consistent difference in mRNA and protein abundance, we found a significant (adjusted p-value=7.4e-3) enrichment for enzymes in central metabolism (e.g. 2-oxoglutarate dehydrogenase, glutamic-oxaloacetic transaminase 1, glutamate dehydrogenase 1, fumarate hydratase FH) (Supplementary Information 5). Moreover, we found that SLC25A13, a mitochondrial aspartate/glutamate transporter, is on average expressed significantly higher and simultaneously is less methylated in glutamine-independent versus -dependent cell lines (Figure 4e).

Together with the observation that the contribution of the aspartate carbon backbone to TCA cycle intermediates was mainly found in malate and fumarate, and to a much less extent in other intermediates of TCA cycle (Figure 4d), our findings are consistent with previous evidence^49^ and altogether point to a potential key role of malate-aspartate shuttle in mediating the ability of cells to use aspartate as a source of glutamate.

### Metabolic *vs* regulatory limitations upon glutamine deprivation

We found that glutamine-independent cell lines exhibited slower growth, increased aspartate consumption and a different regulation of aspartate and glutamine metabolism. However, what mechanism causes glutamine-dependent cells to stop cell growth when glutamine becomes scarce, and whether the same mechanisms are responsible for glutamine-dependency across different cell types remain open questions. To address these questions, we used a high-throughput workflow^64^ to analyze the intracellular dynamics of putatively annotated metabolites in both glutamine-dependent and -independent cell lines at 15 minutes, 24, 48, 72 and 96 hours after switching cells from full medium to media without glutamine. To identify metabolic changes that are specific to glutamine limitation as opposed to a general consequence of growth arrest, we also monitored metabolic changes in response to glucose deprivation (Supplementary Information 6). Of note, in contrast to the limitation of glutamine, upon glucose deprivation none of the 54 cell lines were able to grow (Supplementary Figure 1-2).

We restricted our analysis to 634 ions that were putatively annotated to metabolites with an associated KEGG identifier and initially stratified the major metabolic changes that are common or specific to glutamine and/or glucose deprivation. To that end, we compared the average log_2_ fold-change (FC) of each metabolite across all 54 cell lines during glutamine limitation to the average FC during glucose starvation (Figure 5a-b). We identified 30 and 54 metabolic changes that are specific to either glutamine or glucose deprivation, respectively, and 44 that are common to both starvation conditions (i.e. also with consistent directionality). As expected, most metabolic changes specific to glucose deprivation are involved in sugar and amino sugar metabolism (e.g. UDP-glucose) (Figure 5a). Changes common to glutamine and glucose deprivation are significantly enriched for putative metabolites in arachidonic acid metabolism and central metabolism, especially the TCA cycle, such as lactate, (iso-)citrate, oxoglutarate, fumarate and malate (Figure 5c). Such changes are likely reflecting metabolic effects linked to carbon-starvation and point to the role of glutamine as an anaplerotic carbon source in the TCA cycle. Instead, glutamine-specific changes include several amino acids, such as serine, glycine, valine, threonine, tryptophan, aspartate and methionine, hinting at the key role of glutamine in the global coordination of amino acids metabolism^65^. Notably, four metabolites showed opposing responses to the limitation in glutamine and glucose. Aspartate and AMP showed a strong decrease in glutamine limitation but accumulated in glucose deprivation, while cytidine and NADH showed opposite behavior, i.e. accumulated in glutamine deprivation but decreased in abundance upon glucose starvation, reinforcing a key and specific functional role of aspartate in the homeostatic regulation coordinating glucose and glutamine metabolism.

**Figure 5.**
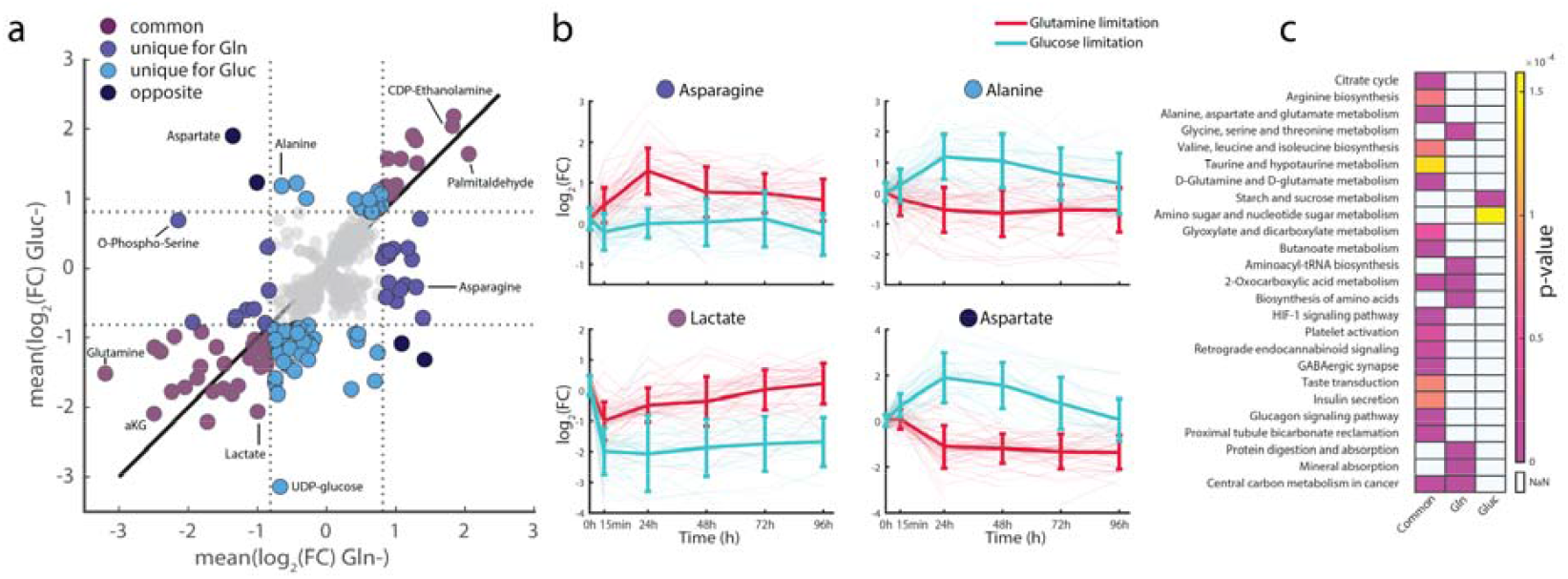
Intracellular dynamics of metabolic adaptation to extracellular glutamine deprivation. **(a)** Metabolic response to glutamine limitation compared to glucose limitation. For each metabolite, the average change to either glutamine or glucose limitation, was calculated across all 54 cell line, respectively. **(b)** Time-dependent fold changes of selected metabolites upon either glutamine limitation (red) or glucose limitation (light blue). The bold line represents the average fold change across all cell lines from the two groups. **(c)** KEGG pathway enrichment of common or condition specific changes in response to glutamine or glucose limitation, as highlighted in panel A. Only pathways with a Bonferroni corrected p-value (1.58e-4) are shown.

Next, we investigated potential differences in the metabolic changes induced by glutamine limitation in glutamine-independent vs. glutamine-dependent cell lines (Figure 6a). We found that 31 metabolites showed a significant difference between the two groups of cell lines in at least one time point after glutamine deprivation (t-test q-value < 0.05 and absolute FC > 0.75) (Figure 5a). Remarkably, the same analysis upon glucose limitation did not yield any significant difference between glutamine-independent and -dependent cell lines (Supplementary Information 6). This suggests that the differential metabolism to bypass glutamine addiction is largely independent from the regulation of glucose metabolism, and most likely points to the essential role of glutamine as a nitrogen donor. While for several of the differential metabolites the changes only gradually emerged over 96 hours, glutamate and N2-Formyl-N1-(5-phospho-D-ribosyl)glycinamide (FGAR) an intermediate of purine metabolism, showed significantly higher abundance in glutamine-independent cell lines already between 24 and 48 hours (Figure 6a). On average, we observed that glutamine-independent cell lines are able to maintain higher levels of glutamate during glutamine deprivation compared to glutamine-dependent cell lines (Figure 6a). This finding is consistent with the observed intrinsic basal metabolic difference between the two groups of cell lines-i.e. glutamine-independent cell lines exhibit a significant conversion of aspartate into glutamate. The contribution from aspartate in the synthesis of glutamate (Figure 3c-d) and slower growth rate (Figure 1d) are likely key factors enabling cells to fulfill glutamine demand for the *de novo* synthesis of biomass precursors (e.g. proteins and nucleotide biosynthesis).

**Figure 6.**
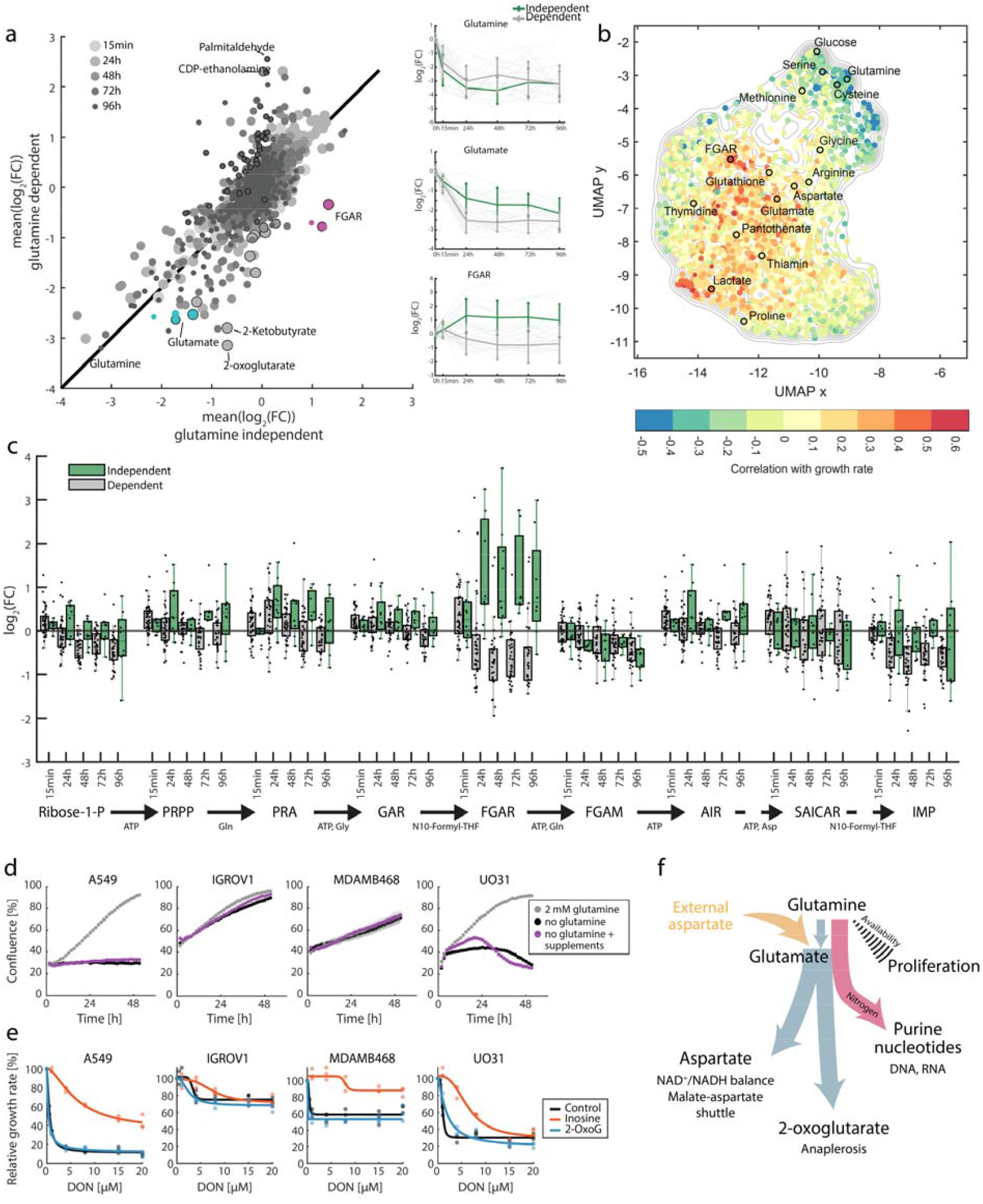
Response of glutamine in/dependent cell lines to extracellular glutamine deprivation. **(a)** Metabolic response to glutamine limitation of glutamine independent cell lines compared to glutamine dependent cell lines. The x-axis shows the average fold change for each metabolite across glutamine independent cell line, the y-axis the average across glutamine dependent cell lines respectively. Dot size indicates the time point. Dots with an outline are metabolites, that show a significant difference between glutamine independent and dependent cell lines (Student’s t-test, multiple testing corrected q-value (Storey) < 0.05). Only ions annotated to KEGG identifiers were considered. **(b)** 2D UMAP projection of exchange fluxes measured for the 54 cell line in RPMI-1640 medium with glutamine. FGAR and selected medium components are highlighted. **(c)** Time-dependent changes in metabolite abundance in the de novo purine synthesis pathway for glutamine independent cell lines (green) and glutamine dependent cell lines (grey). **(d)** Two glutamine dependent (A549 and UO31) and two glutamine independent cell lines (IGROV1 and MDA-MB-468) were grown in medium containing 2 mM glutamine, medium lacking glutamine and medium lacking glutamine but supplemented with IMP, inosine, UMP, uridine, PRPP, AICAR, dihydroorotate, 2-oxoglutarate and GlcNAc (each at 200 µM). Confluence was measured using time-lapse microscopy. Data are shown as mean ± standard deviation (n=3). **(e)** Dose-response curves of cell lines treated with V9302 or DON (glutamine dependent cell lines: A549 and UO31; glutamine independent cell lines: IGROV1 and MDA-MB-468). Cells were supplemented with either 200 µM 2-oxoglutarate (blue line) or 200 µM inosine (orange line). **(f)** Schematic of glutamine and aspartate allocation for catabolic and anabolic reactions.

The second metabolite that exhibits a significant and sustained difference between glutamine-independent and -dependent cells is FGAR, an intermediate in purine *de novo* synthesis. This energetically costly pathway that generates IMP from PRPP is highly regulated ^66 67^ and several metabolic intermediates are known to play key roles in signaling and regulation of the cell cycle progression ^68 69 70^. Two reactions in the pathway consume glutamine as a co-substrate, including phosphoribosylformylglycinamidine synthase (PFAS), the enzyme that converts FGAR into formylglycinamidine-ribonucleotide (FGAM). Possibly because of a slower growth rate, hence lower demand for nucleotide biosynthesis, and the ability to reroute aspartate in glutamate biosynthesis, glutamine-independent cell lines are able to sustain enough flux through the pathway to meet growth requirements and maintain metabolic homeostasis. Remarkably, the enzyme that converts FGAR into formylglycinamidine-ribonucleotide (FGAM) uses glutamine as a co-substrate and on average, glutamine-independent cell lines exhibited an accumulation of FGAR (Figure 6c). Such changes are consistent with glutamine, and not enzyme levels, being the rate-liming factor^71^ for converting FGAR into FGAM. Less obvious are FGAR changes in glutamine-dependent cells. Levels of FGAR and most intermediates in the pathway rapidly decrease upon glutamine deprivation (Figure 6c) pointing to an upstream shutdown of the biosynthetic pathway rather than a mere limitation of glutamine as nitrogen donor.

Therefore, our analysis of metabolic adaptation to glutamine deprivation suggests that purine biosynthesis could be a main factor limiting cell growth *in vitro*. Consistent with this hypothesis, we found that among measured exchange metabolic rates across the 54 cell lines in medium with glutamine, overflow of FGAR strongly correlates with growth rate (fourth most correlated metabolite, Supplementary Information 2) (Figure 6b). Overflow metabolism of metabolic intermediates can report on intracellular flux through the pathway and hint at downstream metabolic bottlenecks^72^ – specifically in this case the conversion of FGAR in FGAM where glutamine is the limiting nitrogen donor. Hence, our metabolomics data suggests a key role of glutamine in allowing faster growth rate by supporting flux in purine biosynthesis as a universal factor limiting growth rate of largely diverse cell types *in vitro*. Remarkably, expression of the pathway enzymes GART and PPAT, but not PFAS for which the metabolic flux is mainly limited by the substrates (glutamine and FGAR) and not enzyme level, were previously shown to be among the most significant prognostic factors of cancer survival in data collected among ∼11‘000 patients with diverse tumor types^73^ (TGCA database, Supplementary Figure 6). This observation hints at similar regulation and growth limiting roles of glutamine and purine biosynthesis *in vivo*.

The next question was whether overcoming metabolic limitations in purine biosynthesis would be sufficient to bypass glutamine dependency. Unexpectedly, we found that neither the supplementation of IMP, nor of downstream products of glutamine metabolism (i.e. UMP, inosine, uridine, dihydroorotate, 2-oxoglutarate, N-acetyl glucosamine, cystine dimethylester, phosphoribosyl pyrophosphate, AICAR) was sufficient to enable glutamine-dependent cell lines to grow in the absence of extracellular glutamine (Figure 6d). Hence, glutamine deprivation might not only act by imposing a metabolic limitation, but also as a signal modulating the basal regulatory state of these cells and causing growth arrest. To test our hypothesis and differentiate the biosynthetic from non-biosynthetic roles of glutamine in mediating cell line-specific dependency on extracellular glutamine, we used the glutamine analog 6-diazo-5-oxo-L-norleucine (DON) to uncouple glutamine biosynthetic and signaling functions. Consistent with our previous observation, glutamine-independent cell lines (i.e. IGROV1, MDAMB468) are resistant to DON, while glutamine-dependent cell lines (i.e. A549, UO31) exhibit high sensitivity (0.5 and 0.97µM IC50, respectively) (Supplementary Information 6). Remarkably, differently from the effects induced by extracellular glutamine limitation, DON-mediated growth inhibition could be rescued by the addition of inosine (Figure 6e, Supplementary Information 6). This finding indicates that besides metabolic limitation of purine biosynthesis, glutamine availability plays a key non-metabolic but signaling role in mediating growth arrest in glutamine-dependent cell lines. Of note, 2-oxoglutarate could not rescue glutamine-dependent cells from DON (Figure 6e, Supplementary Information 6), reinforcing the crucial growth-limiting role of glutamine as a nitrogen donor for purine biosynthesis and not only as an anaplerotic fuel in TCA cycle (Figure 6f).

Altogether, experimental evidence suggests that the signaling cascade mediating growth arrest upon glutamine limitation depends on multiple regulatory events coordinating aspartate uptake, the rewiring of aspartate metabolism into glutamate, slower growth rate, and hence the demand for catabolic and anabolic precursors, including glutamine (Figure 6f).

## Conclusions/Discussion

In recent years, considerable effort to map nutrient dependencies in cancer^12^ has been made. However, a systematic understanding of why some cancer cell types are addicted to specific nutrients while others are not is still missing. Although glutamine addiction was discovered in the 50s^25^, the reasons and mechanisms underlying extracellular glutamine dependency in cancer have been a long standing matter of debate^48,74 32^. Understanding the mechanisms by which extracellular glutamine limitation prevent growth of some cancer cells but not others, and identifying tumors that unequivocally require glutamine as a nutrient, can guide the development of inhibitors of this pathway and the selection of patients that could benefit from such therapeutic approach^75 76 77^. Moreover, a better understanding of the regulation and signaling function of glutamine metabolism can have important implications that extend beyond cancer to immunometabolism, cell differentiation and development^78 59 79^. Here, by comprehensively characterizing the metabolic differences across a panel of 54 cancer cell lines with different levels of dependency on extracellular glutamine, we shed light on common metabolic and regulatory characteristics underlying glutamine in/dependency, that are likely driven by the tissue of origin. We found that purine biosynthesis is a growth limiting factor common to most cell lines, possibly not only *in vitro*, but also *in vivo* where expression of key enzymes in the pathway are strong prognostics factors^13,48,74^. As a result, we showed that the conversion of formylglycinamide ribonucleotide (FGAR) to formylglycinamidine ribonucleotide (FGAM) in the purine biosynthetic pathway is the first metabolic limitation induced by a deficit of glutamine *in vitro*. Remarkably, we found that cells that are independent from extracellular glutamine use extracellular aspartate to synthesize glutamate. Aspartate uptake in combination with a slower growth rate make *de novo* glutamine biosynthesis sufficient to maintain metabolic homeostasis and meet growth biosynthetic requirements. Notably, differential metabolism of glutamine and aspartate seems to be defined by cell identity rather than being an adaptive trait in response to changes in glutamine availability. In fact, similar mechanisms seem to be adopted during cellular differentiation, where undifferentiated human embryonic stem cells consume extracellular aspartate and exhibit a net secretion of glutamine. This pattern is reversed upon differentiation in postmitotic neuronal cells, which consume glutamine and secrete aspartate. However, the inability to fulfill the biosynthetic demand for rapid growth isn’t the only reason conferring glutamine dependency. In fact, even bypassing glutamine induced metabolic limitations by supplementing inosine and/or oxoglutarate, is not sufficient to revert glutamine dependency. On the contrary, we showed that if we expose cells to the glutamine analog DON, even glutamine dependent cell lines are able to grow if inosine is supplemented. This finding highlights the dominant role of glutamine in growth control. In addition, we showed that glutamine in/dependency is likely driven by the tissue of origin and that the growth advantage provided by extracellular glutamine in sustaining high flux in the purine biosynthetic pathway seems a plausible explanation for the fact that most cancer cell lines tested (82%) exhibit glutamine addiction.

By combining the analysis of growth dynamics with the metabolic lifestyles of cancer cells at steady state and upon nutrient deprivation, we propose a general framework to rapidly generate experimentally testable hypotheses on the mechanisms underlying and potentially bypassing nutrient dependencies. A mechanistic understanding of such dependencies can facilitate the rational identification of cancer-specific vulnerabilities. Such mechanistic understanding will enable to translate nutrient dependencies in patient’s derived tumors in strategies that can impact patients’ health and aid in treatment decision making.

## Acknowledgments

We thank Sarah Maria-Fendt, Christian Frezza and all members of the Zampieri group for helpful feedbacks and discussions.

## Author contributions

M.R., K.O., S.D. and M.Z. designed the project. K.O. and S.D. performed the untargeted metabolic profiling. K.O. and M.R. performed tracing metabolomics, data analysis and together with M.B. and L.S. follow-up experiments. M. M. and S.D.G. performed stem cell differentiation experiments. All authors contributed to preparing the manuscript.

## Data and materials availability

All data generated or analyzed during this study are included in this published article as Supplementary Data. A detailed description of all data analysis steps is published in this article in the Methods section and Supplementary Information. Metabolome data and results from data analysis can be found as Supplementary Tables.

## Materials and Methods

### Cell culture

The NCI-60 cancer cell lines were obtained from the National Cancer Institute (NCI, Bethesda, MD, USA). In this study, the 54 adherently growing cell lines were used. The cell lines were expanded from cryopreserved stocks and routinely maintained at 37°C in 5% CO_2_ atmosphere in RPMI-1640 (Biological Industries, cat. no. 01-101-1A) supplemented with 2 mM L-glutamine (Gibco, cat. no. 25030024), 2 g/L D-Glucose (Sigma Aldrich, cat. no. G8644), 5% fetal bovine serum (FBS, Sigma Aldrich, cat. no. F6178) and 1% Penicillin-Streptomycin solution (Gibco, cat. no. 15140122). Prior to metabolomics experiments, the cells were maintained in growth medium containing 5% dialyzed FBS (dFBS, Sigma Aldrich, cat. no. F0392) for at least two passages.

### Cell growth monitoring

Cell growth was monitored by measuring cell confluence, i.e. the surface area of the culture dish covered by cells, using a plate reader for automated bright-field microscopy imaging (TECAN Spark™ 10M). Adherent cell cultures in 96-well plates (Nunc cat. no. 167008, Thermo Scientific) were continuously incubated in the plate reader at 37°C in 5% CO_2_ atmosphere, and images were acquired of each well in 1.5 hours intervals for up to 96 hours. Confluence was estimated from bright-field images using vendor software (TECAN Spark Control). The compilation of growth curves and analysis of growth dynamics was performed in Matlab (MathWorks).

### Glutamine limitation experiments

To monitor changes in growth and metabolome in response to glutamine limitation, we collected cell extracts and culture supernatants at regular intervals during 96 hours exposure of 54 cell lines to either full growth medium (containing 2 mM glutamine) or medium without the addition of glutamine. In each experiment (4 cell lines in parallel), cells were seeded in 96-well plates in triplicates per cell line as described previously^39^. After an initial equilibration and growth phase in full medium, the medium was aspirated from all wells, and was replaced with either full growth medium or medium without the addition of glutamine. Media without glutamine were prepared from glutamine-free RPMI-1640 (Biological Industries, cat. No. 01-101-1A) with 5% dialyzed FBS (Sigma Aldrich, cat. no. F0392), i.e. serum that is devoid of small molecules (10 kDa cutoff) to avoid unintentional supplementation with glutamine. Cell growth was monitored via cell confluence using automated bright-field microscopy (see above). Immediately before, and at 24, 48, 72 and 96 hours after media change, culture supernatants from all wells were transferred to fresh 96-well plates. Adherent cells were extracted with cold organic solvent mixture (40% acetonitrile, 40% methanol and 20% water) with 25 µM phenylhydrazine^80^ as described previously^39^ to generate cell extracts containing intracellular metabolites. Supernatant- and cell extract samples were stored at -80°C until the day of MS analysis. Simultaneously, at each time point, the amount (i.e. confluence) of extracted cells was estimated from replicate cell cultures that were subjected to the same washing step as for cell extraction, but were supplied with PBS instead of extraction solvent. The cultures were then immediately subjected to bright-field microscopy imaging to determine the residual cell confluence, which corresponds to the number of cells present at addition of the extraction solvent.

### Mouse tissue sampling

Prior to tissue collection, RNase- and DNase-free 2.0 mL tubes (Sarstedt, 72.694.406) were pre-filled with 50 mg of ceramic beads (Cayman Chemical, 10402). On the day of tissue collection, three 10-weeks old female FVB mice (line/strain: 1060 - K8-Cre x PIK3CA-H1047R, line/strain ID: 1060, license number: 2924_35058) were euthanized in accordance with institutional animal care and use guidelines. Organs of interest were promptly dissected and sectioned into approximately 50 mg fragments. Each tissue fragment was placed into an individual well of a sterile 24-well plate, which was immediately snap-frozen by immersion in liquid nitrogen to preserve metabolite integrity. Following completion of all dissections, each frozen tissue section was weighed to facilitate accurate downstream quantification and transferred to its corresponding pre-labelled tube containing ceramic beads. An appropriate volume of extraction solvent (40% acetonitrile, 40% methanol and 20% water with 25 µM phenylhydrazine) was then added to each tube to achieve a final tissue-to-solvent concentration of 50 mg/mL and hence allow direct comparison. Finally, samples were stored at -80°C until MS measurements. On the day of MS quantification, tissue samples were first thawed on ice and then homogenized using a Precellys homogenizer equipped with a Cryolys cooling system to preserve metabolite integrity. Dry ice was loaded into the Cryolys detachable cooling unit to maintain a processing temperature of 0°C during grinding. Homogenization was performed in 2 mL tubes 21at 10k RPM for three cycles of 20 seconds each, with an automatic pause of 30 seconds between cycles. Following homogenization, samples were centrifuged at 4°C at maximum speed for 10 minutes to separate the clean supernatant, containing soluble polar metabolites, from beads, tissue debris, and precipitated proteins. The clear supernatant was transferred to a 96-well plate and were subsequently analyzed in randomized order by flow-injection time-of-flight mass spectrometry (FIA-TOF MS, see below).

### Metabolomics profiling of supernatants and estimates of exchange rates

Net uptake and secretion of putatively annotated ions were quantified in 54 adherent cell lines from the NCI-60 panel. Culture supernatants were taken 15 minutes, 24, 48, 72 and 96 hours after supplying cells with fresh growth medium. Samples were analyzed by mass spectrometry-based non-targeted metabolomics to measure the abundance of small molecules. To that end, immediately prior to MS injection, supernatant samples were thawed and diluted 1:20 with deionized water to reduce salt intake and avoid signal saturation. The diluted were subsequently analyzed in randomized order by flow-injection time-of-flight mass spectrometry (FIA-TOF MS, see below). For each of the putatively annotated ion quantified from FIA-TOFMS measurements (as described below), we estimated exchange rates, by first fitting a linear model relating ion intensity to cell numbers over 3 biological replicates (see Figure 2a). We use the robust fitting analysis in Matlab (i.e. function fitlm with RobustFit option) to reduce the influence of outliers in the analysis. For each cell line estimated curve slopes were multiplied by the corresponding cell line growth rates (1/h) and normalized by the cell size correcting factor (a.u.). Cell size characteristic of the 54 cell lines was previously estimated in ^46^ and can be found in the Supplementary Table 2.

### Dynamic metabolic profiling of intracellular metabolic changes

To assess metabolic responses induced by changes in media composition, we measured relative changes in the intracellular abundance of metabolites in cell extracts generated immediately before, and 24, 48, 72 and 96 hours after media change (see above). The cell extract samples were thawed immediately prior to MS analysis, and kept on ice during all subsequent steps. The bottom of each well was scraped with wide-bore pipet tips attached to a multichannel pipet, and the solution was transferred to a fresh 96-well plate with a conical bottom. The plates were sealed and centrifuged at 4k rpm and 4°C to separate cell debris. Cell-free extracts were then transferred to a fresh 96-well plate alongside with quality control- and blank samples (cell-free extraction solvent). The plates were sealed and stored at 4°C in the autosampler until injection and MS analysis (FIA-TOFMS, see below).

Raw MS data (i.e. measured ion abundances) was first annotated by matching the accurate mass of detected ions to known masses of metabolites listed in the Human Metabolome Database (HMDB) and the genome-scale reconstruction of human metabolism (Recon2)^81^ with 3 mDa annotation tolerance (putative annotation). Subsequently, we analyzed ion abundances in quality control samples (12 different pooled cell extract samples) that were measured on each plate, and corrected for batch effects as described previously^46^. To account for differences in the number of cells extracted at each time point, we finally applied a normalization procedure based on linear regression of measured ion abundances at different cell confluences in full medium, as described previously ^46^. The resulting dependency of ion abundance on the number of extracted cells was subsequently used to calculate fold-changes of metabolite abundance in glutamine limitation at each time-point, relative to the expected abundance in full medium at the same cell concentration, as described previously ^46^. Statistical significance (p-value) of each change was assessed by t-test against full medium conditions, and p-values were corrected for multiple hypothesis testing (Benjamini-Hochberg correction at each time point).

### Isotopic tracing with ^13^C-labeled aspartate and glutamine

For the metabolic tracing experiments, A549, UO31 and MDA-MB-468 were seeded in triplicates in 6-well plates (Nunc cat. no. 140675, Thermo Scientific) in RPMI-1640 medium (5% dFBS, 2 mM glutamine, 150 µM aspartate, 1% P/S). After 24 hours, the medium was replaced with labeling medium. RPMI-1640 (5% dFBS, 1% P/S) supplemented with 2mM ^13^C_5_ -glutamine (cat. no. 605166, Sigma Aldrich) was used for glutamine labelling experiments. For ^13^C_4_ -aspartate labeling, RPMI-1640 medium (5% dFBS, 1% P/S) containing 2 mM unlabeled glutamine was supplemented with 150 µM ^15^N-aspartate (cat. no. 332135, Sigma Aldrich). It is noteworthy that RPMI-1640 medium already contains 150 µM aspartate, i.e. in the labeling experiment the total supply of aspartate was 300 µM, 50% of which was ^13^C_4_ -labeled. Samples were collected at two time points (24 and 36 hours after medium replacement) by washing once with 75 mM ammonium bicarbonate (pH 7.4, 37 °C) and immediate addition of extraction solvent (40% methanol, 40% acetonitrile, 20% water, 25 µM phenyl hydrazine, -20°C). The plates were incubated at -20°C for 1 h and then transferred to -80°C until further processing.

Before analysis of the samples, the cells were scraped in order to disrupt the cells and detach them. The supernatant was transferred to a 96 well plate, which was then centrifuged (4°C, 4000 rcf). For FIA-TOFMS analysis, 20 µL of the cell-free extract was transferred to a new plate. For the LC-MS/MS measurement, 150 µl of the cell-free extract was transferred to a new plate and dried in a vacuum centrifuge. The samples were then re-suspended in 50 µL H_2_O for LC-MS/MS analysis.

### Analysis of mRNA, proteome and methylome datasets

Gene expression and methylation datasets were downloaded from Cellminer^62^, while proteome data from ^63^. We identified 3‘039 genes with reported gene expression, protein and methylation levels. For each dataset independently we estimated fold changes and t-test p-values between glutamine dependent and independent cell lines (Supplementary Table S3).

### LC-MS/MS and FIA-TOFMS analysis of ^13^ C-labeled metabolites

LC-MS/MS measurements of metabolites and their isotopologues were carried out on a Waters Acquity UPLC system (Waters Corporation, Milford, MA, USA) coupled to a Thermo TSQ Quantum Ultra (Thermo Scientific), following a method previously published by Buescher *et al*.^82^. The method was adapted to contain the transitions for the metabolites of interest. Because MS/MS transitions for fumarate and the phenyl hydrazine derivative of 2-oxoglutarate (generated during extraction to stabilize the α-keto acid) were not available, the abundance of 2-oxoglutarate isotopologues was assessed from FIA-TOFMS measurements (see description of metabolic profiling).

### Statistical analysis

All data processing and calculations were performed in Matlab 2020a (The Mathworks, Natick). Multiple testing correction was performed using the Storey correction^83^. Where the fraction of true negatives could not be estimated correctly, which is needed for the Storey correction, Benjamini-Hochberg false discovery rate correction was applied^84^. Significance in figures is indicated as follows: * p < 0.05, ** p < 0.01, *** p < 0.001, **** p < 0.0001, ns: no significance.

### Virus production

Lentivirus was produced by transfection of lentiviral backbones containing the indicated transgenes together with third-generation packaging plasmids into HEK293 cells following the Trono laboratory protocol^85^. Virus was concentrated from culture supernatant by ultra-centrifugation (23k rpm, 2 hours, 4°C) and either stored at -80°C or used directly for transduction.

### Generation of human neurons

Human neuron generation by transcription factor overexpression has been described previously^86^. In brief, on day -1 human stem cells (H1; WiCell Research Institute (WA)) were treated with Accutase (Gibco) and plated on Matrigel (Corning)-coated 6-well plates in mTeSR Plus containing 10 µM Y-27632 (Axon Medchem). While plating, lentivirus encoding doxycycline-inducible Ngn2 (Addgene Plasmid #52047) was used to transduce the cells. On day 0, Doxycycline (2 mg/ml, Sigma) was added to induce Ngn2 expression and retained in the medium throughout the experiment. On day 1, cells were treated with Accutase (Gibco) and plated on Matrigel (Corning)-coated 6-well plates in DMEM/F12 containing N2-supplement, non-essential amino acids (all from Invitrogen), human BDNF (10 ng/ml, PeproTech), human NT-3 (10 ng/ml, PeproTech), and mouse Laminin-1 (0.2 mg/ml, Invitrogen) containing 10 µM Y-27632 (Axon Medchem) and puromycin (2 µg/ml c.f.) was added for 3 days to select infected cells. On day 4, neuronal cells were harvested for metabolomics or further matured by changing into Neurobasal-A medium supplemented with B27/glutamax (Invitrogen) containing BDNF, NT3, Laminin-1, and Ara-C (2 µM, Sigma) with media changes at day 8 until cells were harvested for metabolomics on day 10. All cells were grown at 37°C and 5% CO_2_.

