## Supplementary Figures for "The glutamine-aspartate metabolic tradeoff between fast growth and metabolic flexibility"

**Affiliations:**

**Supplementary Materials**

### 1 **Supplementary Data**

*Available for download as Excel files from Journal website.*

**Supplementary Information 1.** Continuous monitoring of confluence in RPMI media with and without glutamine or glucose in 54 NCI-60 cell lines.

**Supplementary Information 2.** Estimates of relative uptake/secretion rates and dynamic fold change analysis of intracellular metabolites upon glutamine or glucose deprivation in 54 NCI-60 cell lines.

**Supplementary Information 3.** Mass spectrometry data monitoring the incorporation of the isotopic labels from ^13^C_5_-glutamine and ^13^C_4_--aspartate in A549, MDA-MB-468, and UO31 cell lines.

**Supplementary Information 4.** Metabolome profiles of 12 mouse organs together with intracellular and extracellular metabolic changes in human embryonic stem cells (hESC) cells during transcription factor-mediated differentiation into excitatory induced neurons.

**Supplementary Information 5.** Analyses of previously published transcriptomic, proteomics, drug IC50 and methylome data.

**Supplementary Information 6.** Dynamic metabolome changes upon limitation of glucose or glutamine in 54 adherent cell lines.

### 2 Supplementary Figures

**Supplementary Figure 1.** Continuous monitoring of confluence in RPMI media with and without glutamine or glucose in 54 NCI-60 cell lines.

**Supplementary Figure 2.** Estimates of relative growth index in RPMI media without glutamine or glucose

**Supplementary Figure 3.** Mouse organ atlas and fold-change volcano plots of 2170 putatively annotated intracellular metabolites after neuronal induction.

**Supplementary Figure 4**. Continuous monitoring of confluence in RPMI media lacking glutamine with different concentrations of aspartate dimethyl ester.

**Supplementary Figure 5.** mRNA levels of GLUL and GLS in glutamine (in)dependent cell lines.

**Supplementary Figure 6**. hazard ratios (HRs) and P-values for gene expression levels of all metabolic enzymes (1126 enzymes) in cancer cohort studies

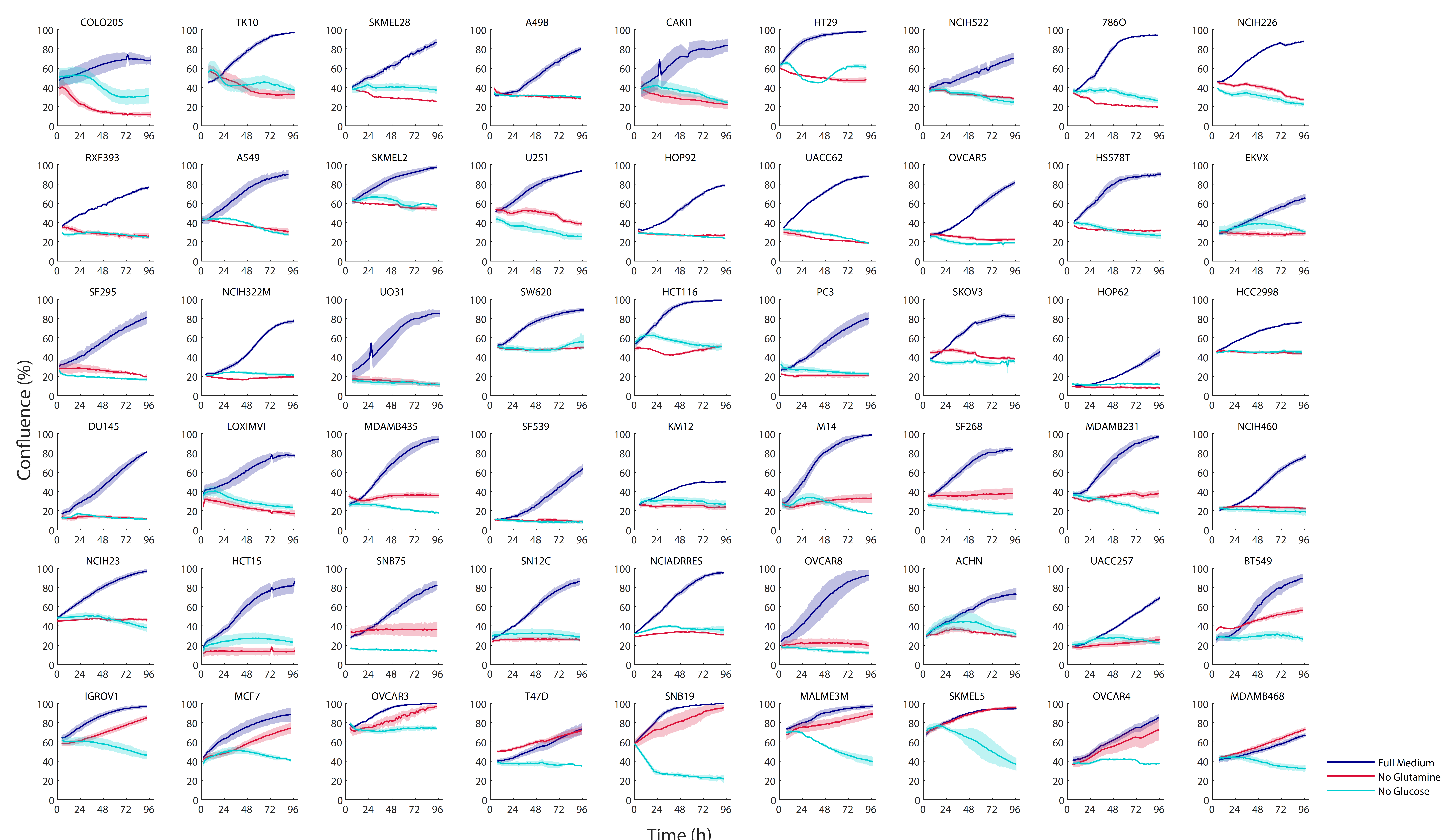
**Supplementary Figure 1:** Continuous monitoring of confluence in RPMI media with and without glutamine or glucose in 54 NCI-60 cell lines.

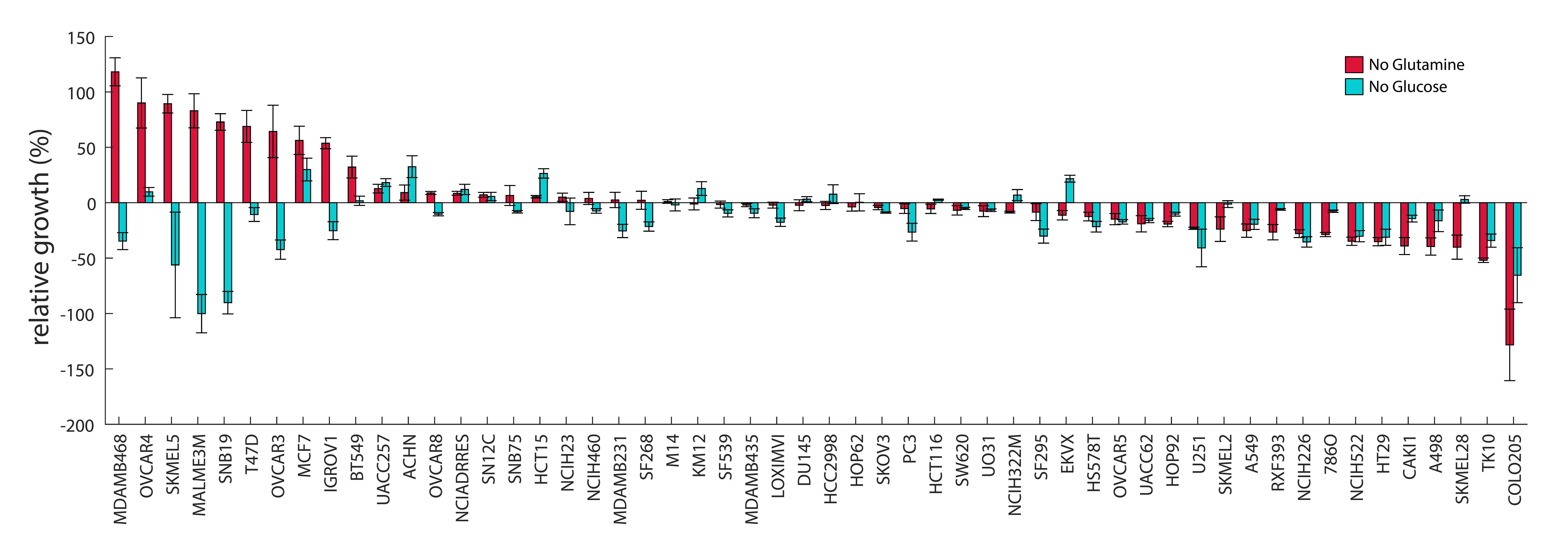

**Supplementary Figure 2.** Estimates of relative growth in RPMI media without glutamine or glucose

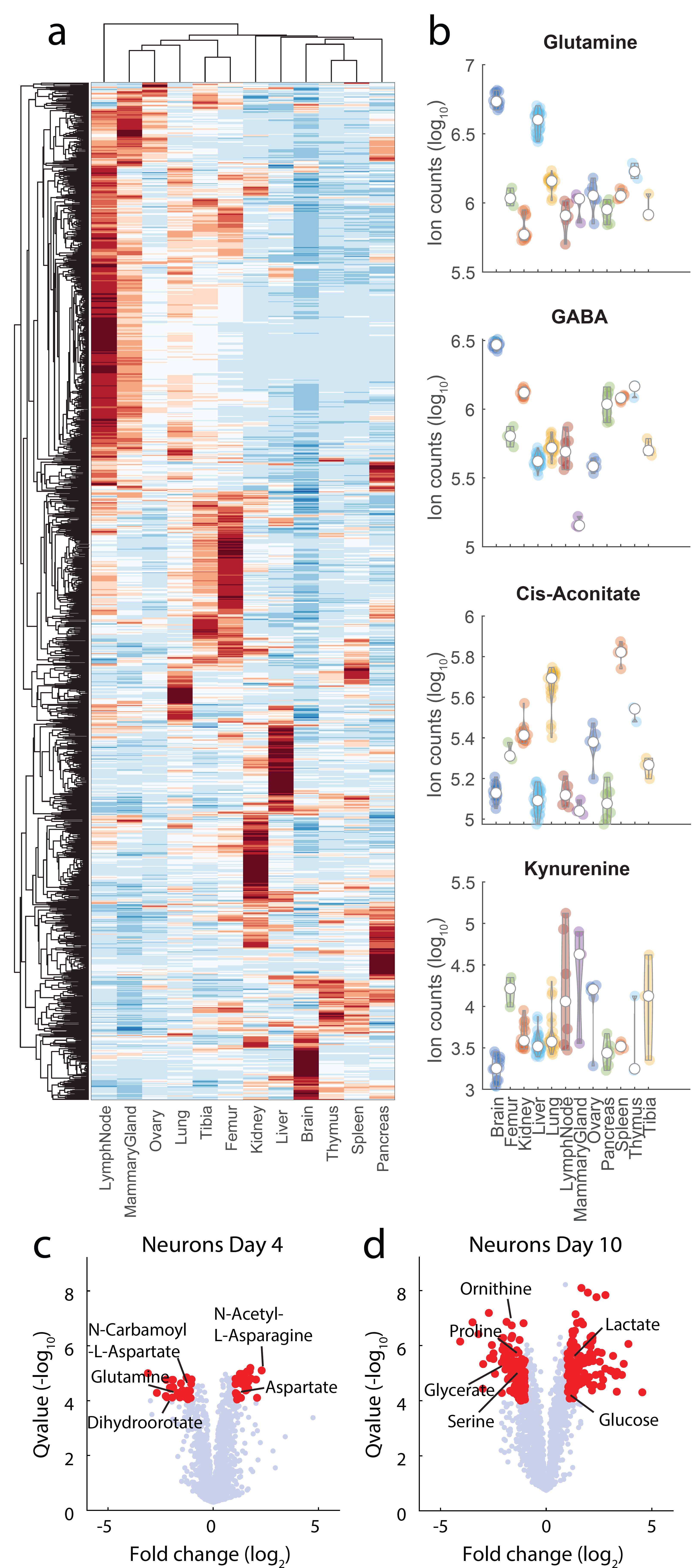

**Supplementary Figure 3. (a)** Clustergram of z-scored normalized levels for XYZ putatively annotated metabolites in 12 organs sampled at multiple sites from 3 FVB mice. **(b)** Raw intensity counts for 4 selected metabolite examples. **(c-d)** Fold-change volcano plots of 2170 putatively annotated intracellular metabolites 4 **(c)** and 10 **(d)** days after neuronal induction. Selected annotated metabolites exhibiting significant (q-value<1e-4 and log2 fold-change >=1) changes during differentiation are highlighted in red. We observed initial changes in the concentrations of intermediates of aspartate metabolism (e.g. aspartate, N-carbamoyl-aspartate), followed by widespread metabolic changes in central metabolism, including significant changes of CoA, Acetyl-CoA and several intermediates of TCA cycle (e.g. (iso)citrate, oxaloacetate).

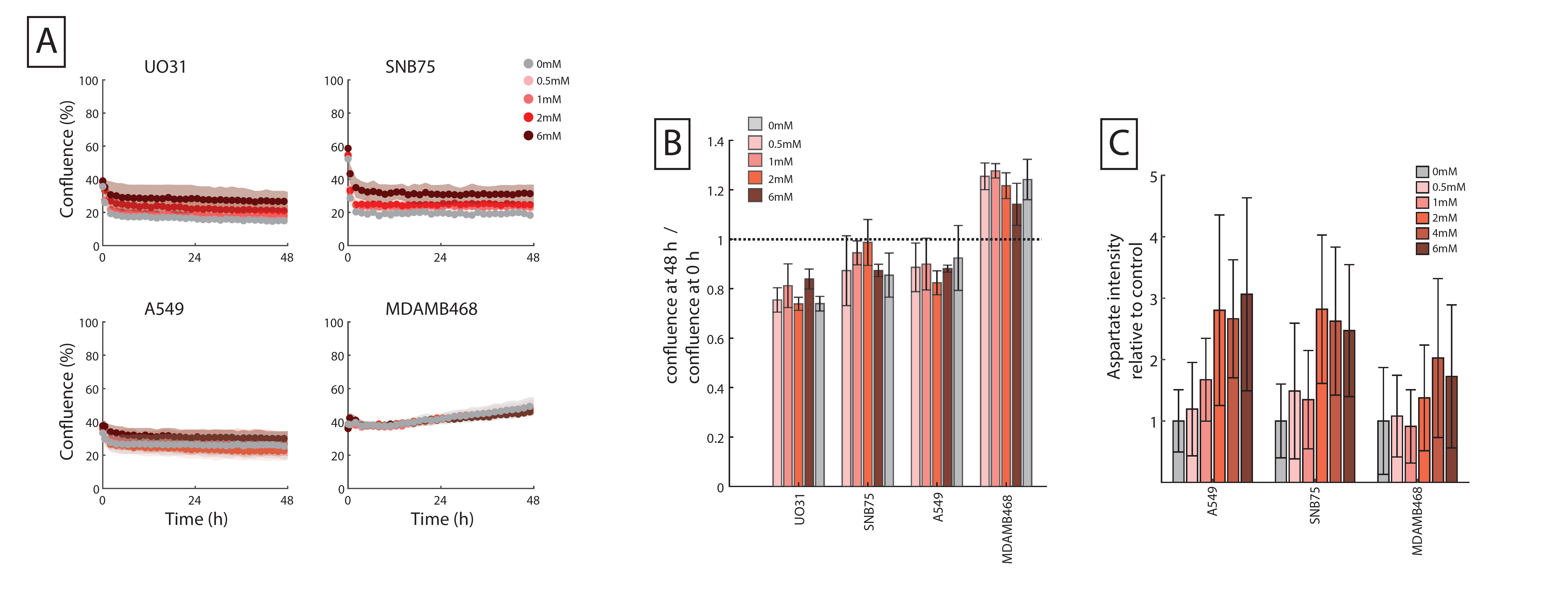

**Supplementary Figure 4. (a)** Continuous monitoring of confluence in RPMI media lacking glutamine with different concentrations of aspartate dimethyl ester of 4 cell lines: three glutamine dependent (A549, UO31 and SNB75) and one glutamine independent cell line (MDA-MB-468, in green). **(b)** The confluence 48 h after glutamine deprivation is divided by the confluence before glutamine deprivation and reported as a ratio (i.e. relative confluence). C) Intracellular lelves of aspartate fpr 3 cell lines grown in in RPMI medium with glutamine and different coenctrations of aspartate dimethyl ester.

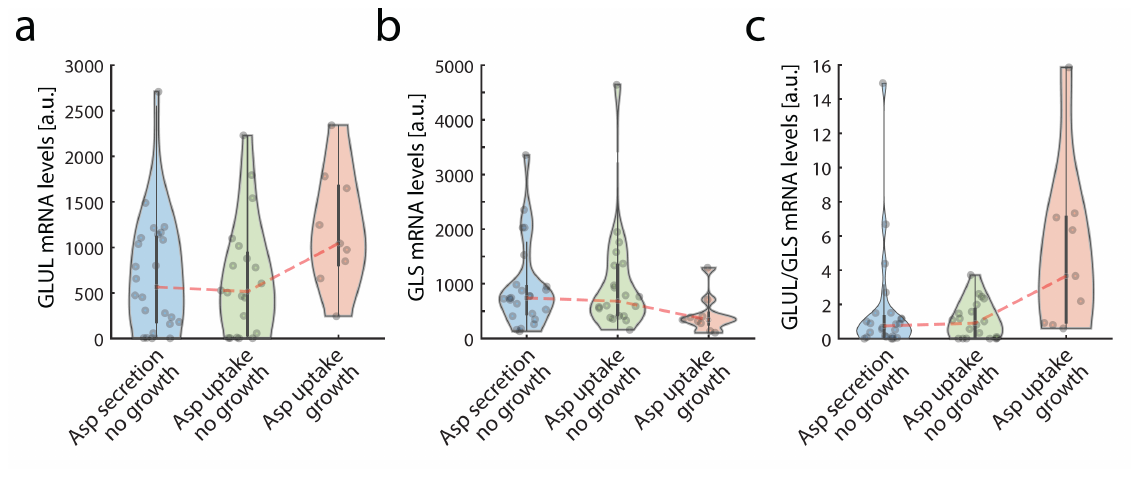

**Supplementary Figure 5. (a)** Previously published^1^ mRNA levels of GLUL, **(b)** GLS and the **(c)** ratio between GLU and GLS for the 54 cell lines grouped by glutamine independent (red), glutamine dependent with a net uptake (green) or a net secretion of aspartate (blue).

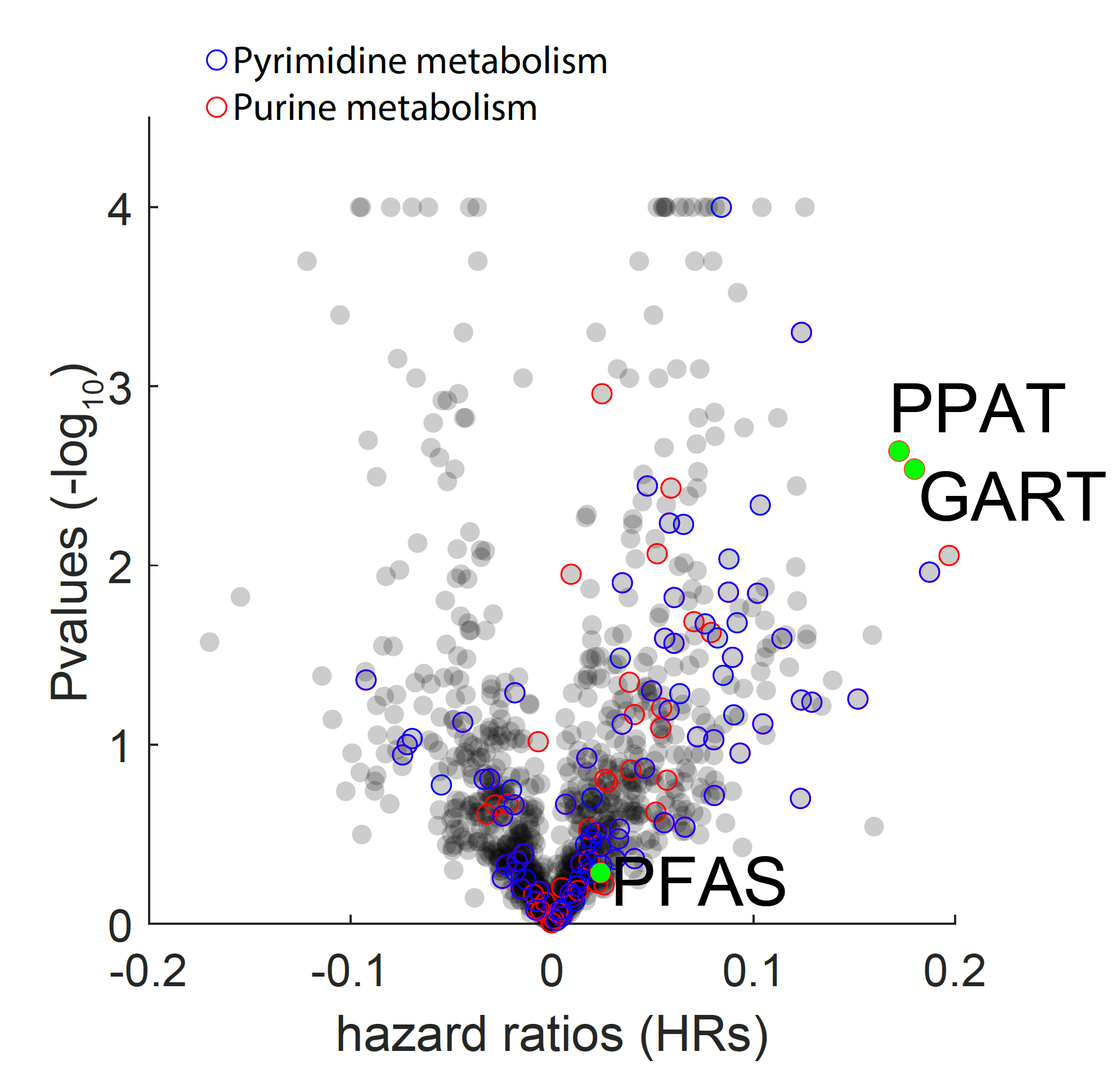

**Supplementary Figure 6**. In this volcano plot we reported previously published^2^ integrated hazard ratios (HRs) and P-values for gene expression levels of all metabolic enzymes (1126 enzymes) in cancer cohort studies. Enzymes of the purine and pyrimidine biosynthetic pathways are shown in red and blue, respectively.
